# A simple, cost-effective, and robust method for rRNA depletion in RNA-sequencing studies

**DOI:** 10.1101/2020.01.06.896837

**Authors:** Peter H. Culviner, Chantal K. Guegler, Michael T. Laub

## Abstract

The profiling of gene expression by RNA-sequencing (RNA-seq) has enabled powerful studies of global transcriptional patterns in all organisms, including bacteria. Because the vast majority of RNA in bacteria is ribosomal RNA (rRNA), it is standard practice to deplete the rRNA from a total RNA sample such that the reads in an RNA-seq experiment derive predominantly from mRNA. One of the most commonly used commercial kits for rRNA depletion, the Ribo-Zero kit from Illumina, was recently discontinued. Here, we report the development a simple, cost-effective, and robust method for depleting rRNA that can be easily implemented by any lab or facility. We first developed an algorithm for designing biotinylated oligonucleotides that will hybridize tightly and specifically to the 23S, 16S, and 5S rRNAs from any species of interest. Precipitation of these oligonucleotides bound to rRNA by magnetic streptavidin beads then depletes rRNA from a complex, total RNA sample such that ~75-80% of reads in a typical RNA-seq experiment derive from mRNA. Importantly, we demonstrate a high correlation of RNA abundance or fold-change measurements in RNA-seq experiments between our method and the previously available Ribo-Zero kit. Complete details on the methodology are provided, including open-source software for designing oligonucleotides optimized for any bacterial species or metagenomic sample of interest.

**Importance:** The ability to examine global patterns of gene expression in microbes through RNA-sequencing has fundamentally transformed microbiology. However, RNA-seq depends critically on the removal of ribosomal RNA from total RNA samples. Otherwise, rRNA would comprise upwards of 90% of the reads in a typical RNA-seq experiment, limiting the reads coming from messenger RNA or requiring high total read depth. A commonly used, kit for rRNA subtraction from Illumina was recently discontinued. Here, we report the development of a ‘do-it-yourself’ kit for rapid, cost-effective, and robust depletion of rRNA from total RNA. We present an algorithm for designing biotinylated oligonucleotides that will hybridize to the rRNAs from a target set of species. We then demonstrate that the designed oligos enable sufficient rRNA depletion to produce RNA-seq data with 75-80% of reads comming from mRNA. The methodology presented should enable RNA-seq studies on any species or metagenomic sample of interest.

## Introduction

RNA-sequencing (RNA-seq) is a common and powerful approach for interrogating global patterns of gene expression in all organisms, including bacteria(1–3). In most RNA-seq studies, it is desirable to eliminate rRNAs so that as many reads as possible come from mRNAs(4). For most eukaryotes, the majority of mRNAs are polyadenylated, enabling their selective isolation and subsequent sequencing(5, 6). In contrast, bacteria do not typically poly-adenylate their mRNAs, and rRNA comprises 80% or more of the total RNA harvested from a given sample(7). To enrich for mRNA in RNA-seq samples, a general strategy involves the depletion of rRNAs by subtractive hybridization(8–10). This approach was at the heart of commercially available kits such as Ribo-Zero from Illumina, leading to RNA-seq data in which ~80-90% of the reads map to mRNAs. Despite the popularity and efficacy of Ribo-Zero, this kit was recently discontinued by the manufacturer.

Here, we report an easily implemented, scalable, and broadly applicable do-it-yourself (DIY) rRNA depletion kit. Our kit relies on the physical depletion of rRNA from a complex RNA mixture using biotinylated oligonucleotides (oligos) specific to 5S, 16S, and 23S rRNA. We focus primarily on the development of oligos that will enable depletion of rRNA from any one of eight different, commonly studied bacteria. However, we also present an algorithm for customizing the subtractive oligos, and the open-source software developed here can be used to design oligonucleotides for the depletion of rRNA from any user-defined set of species. Our results indicate that the kit we developed enables the facile depletion of rRNA from total RNA samples such that ~70-80% of reads in RNA-seq map to mRNAs. We further demonstrate that our kit produces RNA-seq data showing high correspondence to that produced using Ribo-Zero kit. Additionally, our kit has a reduced cost of only ~$10 per sample to deplete rRNA from 1 μg of total RNA. We anticipate that this rRNA-depletion strategy will benefit the entire bacterial community by enabling low-cost transcriptomics with a similar workflow to the previously available Ribo-Zero kit.

## Results

To efficiently and inexpensively deplete rRNA from total RNA from multiple organisms, we developed an algorithm to design DNA oligonucleotides capable of hybridizing to rRNA from multiple species simultaneously. We reasoned that each rRNA should be bound by multiple oligos across the length of the rRNA, in case a given site is hidden by structure or is not available due to partial fragmentation during RNA extraction or processing. Further, we decided that oligos should be as short as possible to reduce synthesis cost and decrease the likelihood of spurious binding and accidental depletion of mRNA. To find potential binding sites, we aligned the 16S and 23S rRNA sequences from a set of eight commonly studied bacteria, including several major pathogens and model organisms (Figure 1A, S1A). These sequences were divergent enough that we could not design an oligo based on the rRNA sequence of a single species and expect it to bind other the rRNA from other species effectively. Thus, we designed an algorithm to optimize the sequence of oligonucleotides, enabling them to hybridize to rRNA from multiple species. To find these oligos, we focused on ungapped regions of the alignment and chose a large number of sites as candidates (Figure 1B, S1B). For nucleotide positions that were completely conserved among the eight species, the conserved nucleotide was selected. For positions that were only partially conserved, a nucleotide was chosen at random such that it would match a nucleotide found in some but not all of the rRNAs.

**Figure 1.**
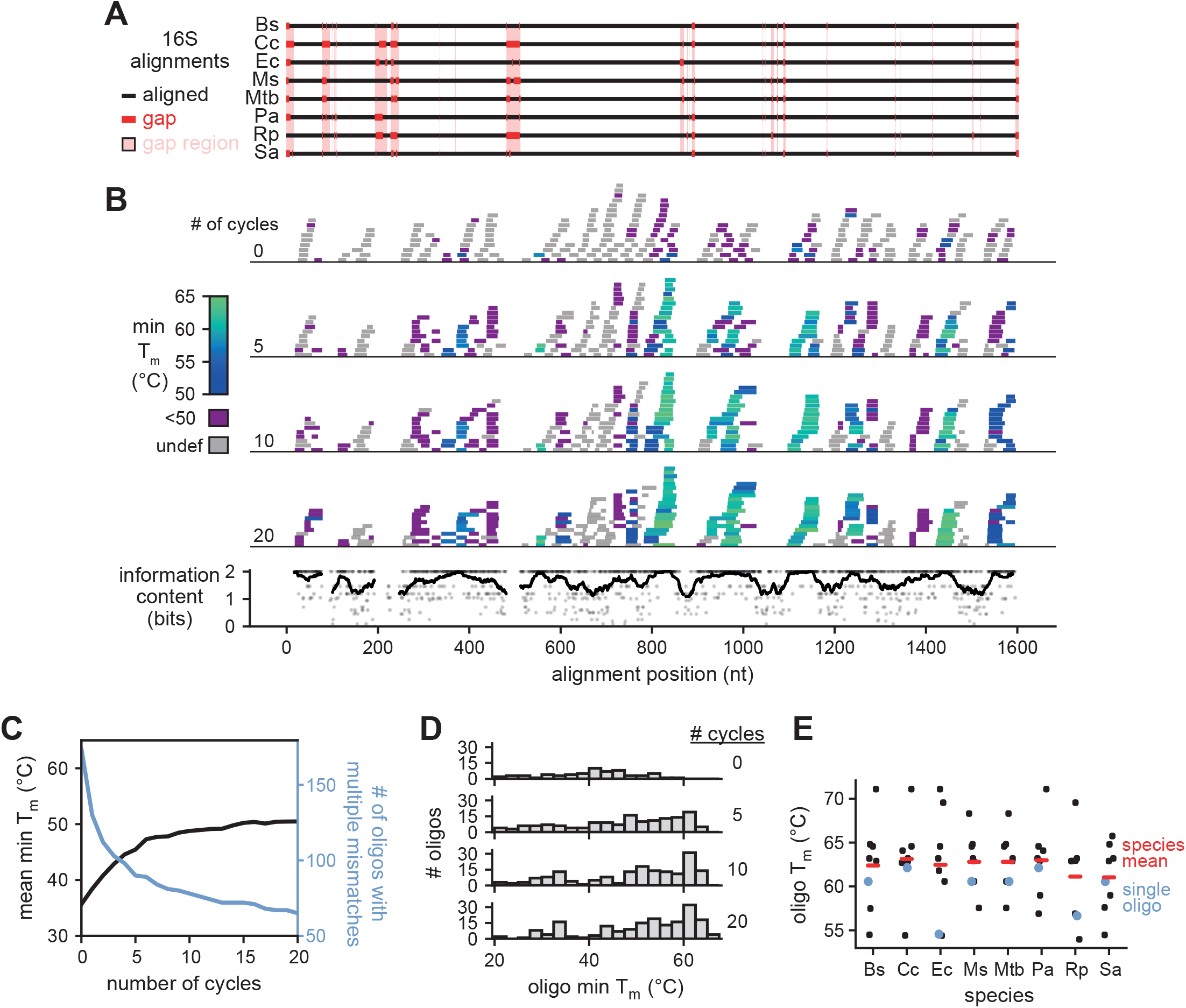
Oligonucleotide selection for 16S rRNA. (A) Alignment of 16S sequences from 8 bacterial species (Ec = *E. coli*; Pa = *P. aeruginosa*; Rp =*parkeri*; Cc = *C. crescentus*; Bs = *B. subtilis*; Ms = *M. smegmatis*; Mtb = *M. tuberculosis*; Sa =*aureus*). Alignment gaps are shown as red lines in the particular species of the gap. Regions with a gap in any species are highlighted in pink; these regions were not considered when designing oligos. (B) The position, length, and minimum T_m_ of all oligos plotted against the 16S alignment after the indicated number of optimization cycles (top). The information content at each nucleotide position of aligned regions is also shown (bottom, points). To highlight conserved regions, a sliding average information content is also plotted (bottom, line). (C) Oligo T_m_ statistics after multiple cycles of the T_m_ optimization algorithm. For each oligo (n = 250), we calculated the minimum T_m_ across the 8 species considered and then plotted the mean of this value across all oligos (black). The T_m_ cannot be accurately estimated for oligos with multiple sequential mismatches; the number of oligos with an undefined T_m_ is also plotted (blue). (D) Histograms of minimum T_m_ for oligos at the indicated number of optimization cycles. Data were generated as in (C), but oligo T_m_ minima were used to generate histograms rather than taking the mean across all oligos. Oligos with an undefined T_m_ were not included in the histograms. (E) Distribution of T_m_ values for each 16S-targeting oligo (n = 8) for each individual species indicated. The mean T_m_ of oligos for each species is also shown (red lines). Note that the same oligos are used for each species, but because of 16S sequence variability, the T_m_ can vary, as illustrated for one particular oligo (blue).

We then performed an iterative process to sample alternate sequences and binding locations for each oligo, while biasing the selection toward sequences that tightly bind rRNA from the eight species we had selected. To do this, for each oligo we generated *in silico* a set of mutated oligonucleotides that varied from the original sequence by either extending, shrinking, or shifting the binding site, or by mutating a single nucleotide of the oligonucleotide to match a different species’ rRNA. From this set of mutated oligonucleotides, the algorithm effectively replaced each old oligonucleotide with a new one, favoring those close to a target minimum T_m_ of 62.5 °C. This T_m_ was chosen to achieve tight binding across all species while preventing selection of excessively long oligonucleotides. With more cycles of optimization, the average minimum T_m_ approaches the target T_m_ (Figure 1C, S1C). Notably, many oligonucleotides, particularly those with poorly conserved start locations, were not able to reach the target T_m_, though the number that did increased with additional cycles (Figure 1D, S1D). After 15-20 cycles, the oligonucleotides had converged on highly conserved regions of the rRNAs (Figure 1B-D, S1B-D). After 100 cycles of optimization, we selected 8 and 9 non-overlapping oligonucleotides for the 16S and 23S rRNA, respectively, with an average length of 30 nucleotides. These 17 oligos are predicted to hybridize to rRNA from all eight species included in the initial design. Although the T_m_ for individual oligos varies across species, the mean T_m_ for the oligo set as a whole was similar (Figure 1E, S1E).

We also applied our algorithm to the 5S rRNA from the 8 species considered. However, because the 5S rRNA is both shorter and more poorly conserved than 16S and 23S rRNA, we were unable to find oligos that are predicted to effectively hybridize to the 5S rRNA from all eight species. Therefore, we ran the algorithm against individual 5S rRNAs and hand-selected two oligos specific to the 5S from each species. In addition, we found that the algorithm was unable to find oligos mapping near the 5’- and 3’-ends of the 23S due to its low conversation among species. To improve binding to these regions, we also identified two oligos that were specific to either Gram-positive or Gram-negative members of our target set of species. Thus, our final set of depletion oligos for a given organism includes 21 total oligos: 17 common oligos targeting 16S and 23S rRNA, 2 oligos that target 23S rRNA in a Gram positive- or Gram-negative-specific manner, and 2 species-specific 5S targeting oligos (Table S1; oligos each contain a 5’-biotin modification).

We then sought to determine whether our oligo libraries could effectively deplete bacterial rRNA. To deplete rRNAs from a total RNA sample, we incubated biotinylated versions of the 21 designed oligos with total RNA. Samples were then combined with magnetic streptavidin beads to precipitate the oligos bound to rRNAs, followed by isolation of the supernatant, which should be heavily enriched for mRNA (Figure 2A). We extracted total RNA from exponentially growing cultures of common lab strains of *E. coli*, *B. subtilis,* and *C. crescentus* and performed a single round of rRNA depletion. For all three species, incubation with the 21 depletion oligos substantially decreased the intensity of rRNA signal on a polyacrylamide gel, while tRNA and ncRNA were generally unaffected (Figure 2B). Moreover, this depletion was modular, as incubation of *E. coli* total RNA with probes targeting only 16S, 23S, or 5S rRNA resulted in selective depletion of the band corresponding to a given targeted rRNA (Figure 2B, left).

**Figure 2.**
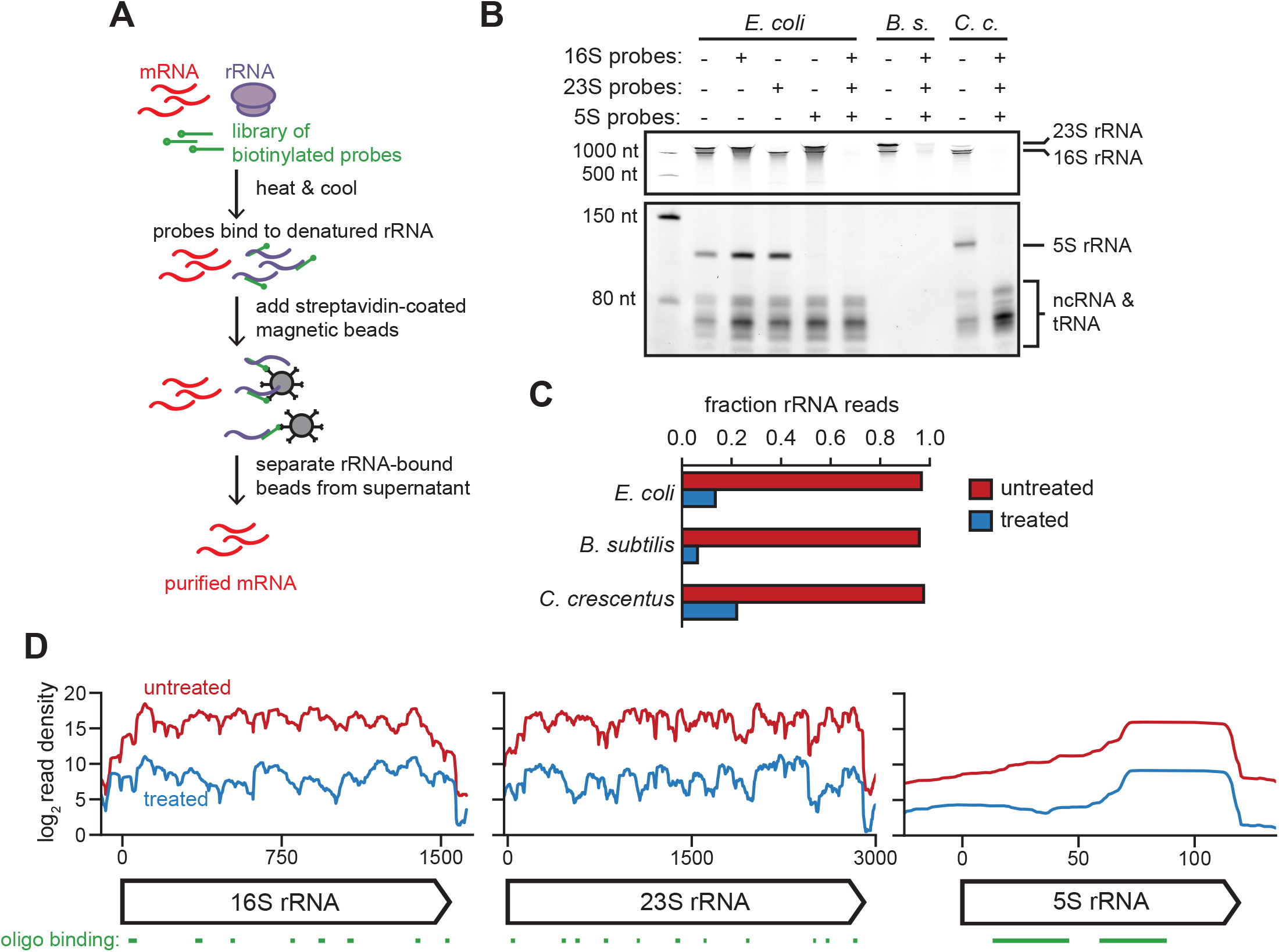
rRNA depletion by oligonucleotide-based hybridization. (A) Cartoon of the rRNA depletion process. (B) Polyacylamide gel showing total RNA from *E. coli*, *B. subtilis*, and *C. crescentus* pre- and post-rRNA depletion using indicated probe sets. The first lane is a ladder. Approximate positions of abundant RNAs, including rRNAs, is indicated on the right. Note that a lower contrast is shown for the top portion of the gel to resolve 16S and 23S bands. (C) Fraction of total reads aligning to rRNA for rRNA-undepleted and -depleted samples of *E. coli*, *B. subtilis*, and *C. crescentus* total RNA. (D) Summed read counts across the *E. coli* 16S, 23S, and 5S rRNAs pre-(red) and post-(blue) depletion. The positions of oligos used for depletion are shown below.

To quantify how well our method depleted rRNA, we performed RNA-seq on the three RNA samples pre- and post-rRNA depletion for *E. coli*, *B. subtilis*, and *C. crescentus*. We then calculated the fraction of reads mapping to rRNA loci in each case (Figure 2C). The fraction mapping to rRNA decreased following depletion from >95% to 13%, 6%, and 22%, respectively. To determine whether there was any bias for depletion of certain regions of the rRNAs, we compared read counts at each nucleotide position pre- and post-depletion in each *E. coli* rRNA (Figure 2D). For the 16S, 23S, and 5S rRNAs, read density was relatively uniform but lower, following depletion, indicating that no particular region of the rRNAs (e.g. regions prone to high structure or partial degradation, preventing effective depletion) was over-represented in our rRNA reads.

Many RNA-seq studies are aimed at detecting significant differences in the expression of mRNA in different strains or across different perturbations. To ensure that our depletion technique did not affect the measurement of expression changes (e.g. through unintended depletion of particular mRNAs), we treated *E. coli* cells with either rifampicin or chloramphenicol for 5 minutes and compared fold changes measured from libraries generated using our depletion strategy to those generated using the previously available commercial kit Ribo-Zero (Illumina). For each depletion method, we calculated the log_2_ fold-change in read counts in coding regions following antibiotic treatment compared to a negative control (Figure 3A-B). For both rifampicin and chloramphenicol treatment, the correlation in log_2_ fold-change per coding region between the two rRNA depletion strategies was high (R^2^ = 0.98 and 0.97 for rifampicin and chloramphenicol, respectively) across a wide range of changes in gene expression. These results indicate that our method should provide similar results to the Ribo-Zero kit for studies measuring changes in gene expression.

**Figure 3.**
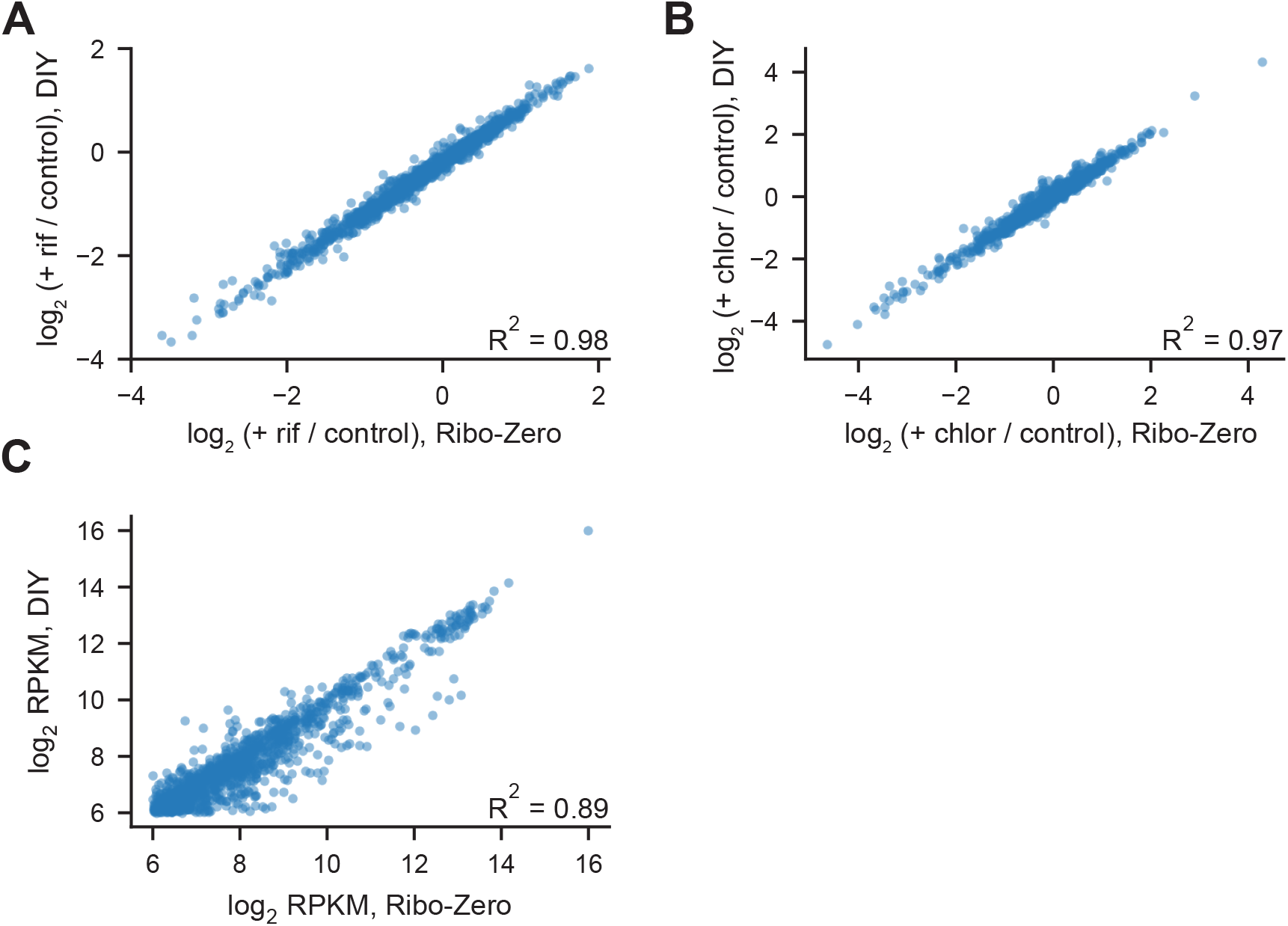
Our rRNA depletion strategy performs comparably to Ribo-Zero for RNA-seq. (A) Scatterplot showing correlation between log_2_ fold changes for *E. coli* coding regions following rifampicin treatment, comparing rRNA depletion via Ribo-Zero with our depletion strategy. Fold changes were calculated as the ratio of RPKM between rifampicin treated and untreated samples. All coding regions with at least 64 RPKM in both untreated samples (n = 1294) were considered in the analysis. (B) Scatterplot showing correlation between log_2_ fold changes for *E. coli* coding regions following chloramphenicol treatment, comparing rRNA depletion via Ribo-Zero with our depletion strategy. Fold changes were calculated as the ratio of RPKM between chloramphenicol-treated and untreated samples. All coding regions with at least 64 RPKM in both untreated samples (n = 1294) were considered in the analysis. (C) Scatterplot showing correlation between read counts (RPKM) for *E. coli* coding regions treated with Ribo-Zero and our do-it-yourself (DIY) depletion strategy. All coding regions with at least 64 RPKM in both samples (n = 1294) were considered in the analysis.

Importantly, we also determined if our kit differentially depleted particular mRNAs compared to the Ribo-Zero kit. To do this, we directly compared the RPKM (reads per kilobase per million) values for well-expressed coding regions from a library prepared using our depletion strategy and one prepared using Ribo-Zero (Figure 3C). Overall, there was a high correlation between the two depletion methods (R^2^ = 0.89). However, there were a few outliers. We first identified genes more than two standard deviations away from a log-linear fit of RPKM in a comparison of RNA-seq data generated using our method and Ribo-Zero (Figure S2A). To ensure that fold-change in expression could also be accurately calculated for these outliers, we returned to the data generated following antibiotic treatment (Figure 3A-B). For the outlier genes, the change in expression following treatment with rifampicin or chloramphenicol was still highly correlated (Figure S2B-C; R^2^ = 0.95 and 0.90, respectively). For a more stringent cut-off, we also hand-selected 11 of the most highly expressed outliers that were more significantly depleted by our method than Ribo-Zero (Figure S2D). Again, the changes in expression calculated following treatment with rifampicin or chloramphenicol were highly correlated (Figure S2E-F; R^2^ = 0.96 and 0.89 for rifampicin and chloramphenicol, respectively). Thus, results obtained based on our oligonucleotide hybridization approach are highly comparable to those generated with the previously-available Ribo-Zero kit.

Finally, we compared the RPKM values for well-expressed coding regions between libraries prepared using our depletion strategy and libraries from total RNA (no rRNA depletion) from *E. coli*, *B. subtilis*, and *C. crescentus* (Figure S3A). Depleted libraries for each species showed similar correlations of R^2^ = 0.76, 0.71, and 0.66 for *E. coli*, *B. subtilis*, and *C. crescentus*, respectively (Figure S3A). Though this analysis is complicated by the relatively few reads mapping to mRNA in undepleted samples (Figure 2C), these results confirm that our method is an effective strategy for depleting rRNAs while maintaining transcriptome composition across multiple species. Taken all together, we conclude that our DIY method provides a broadly applicable, customizable, and cost-effective technique for determining changes in bacterial gene expression patterns in a wide range of organisms and experimental contexts.

## Discussion

We have developed a simple, fast, easy-to-implement, and cost-effective method for efficiently depleting rRNA from complex, total RNA samples. For three different species, *E. coli*, *B. subtilis*, and *C. crescentus*, we demonstrated robust depletion of 23S, 16S, and 5S rRNAs in a single step such that ~70-90% of reads in RNA-seq arise from non-rRNA sources. This level of mRNA enrichment is sufficient for most RNA-seq studies. Our method showed relatively uniform depletion of rRNAs and minimal, unwanted ‘off-targeting’ of mRNAs. Additionally, expression changes measured using our method correlated very strongly (R^2^ = 0.98, 0.97; Figure 3A-B) to those measured using the previously available Ribo-Zero kit. This strong correlation both validates our method and ensures that data generated via either method can be safely compared or combined.

Another method has recently been developed as an alternative to the discontinued Ribo-Zero kit(11). This method is based on hybridization of DNA oligonucleotides to rRNAs followed by digestion with RNase H, which recognizes DNA:RNA hybrids. This method also enabled robust rRNA depletion, although a direct comparison of RNA-seq counts per gene generated using this method and Ribo-Zero was not reported. Additionally, this alternative method requires extended (~60 min) incubations with an RNase, albeit one that should be specific to DNA-RNA hybrids, whereas ours involves only hybridization and a precipitation step.

The set of biotinylated oligonucleotides tested here were designed to deplete the rRNA from a set of 8 selected organisms. These organisms span a large phylogenetic range so these oligonucleotides are likely broadly applicable to different bacterial species or even metagenomic samples. However, the set of oligonucleotides can also be easily optimized for a different species or set of species using the open-source software developed here and available on Github. As noted, because the 5S rRNA is shorter and less conserved, probes specific to the 5S from a given species must typically be designed. However, the 5S rRNA does not yield nearly as many reads in RNA-seq data for a total RNA sample and may not require depletion for all studies.

In sum, the rRNA depletion methodology developed here should facilitate RNA-seq studies for any bacterium of interest. Notably, our method is also substantially cheaper than the Ribo-Zero kit. The cost of our method is ~$10 per reaction to deplete 1 μg of total RNA (see Methods) compared to ~$80 per reaction for Ribo-Zero. The cost for our approach stems primarily from the magnetic streptavidin beads used to precipitate the biotinylated oligonucleotides bound to rRNA. Further optimization of the method reported here could likely reduce the cost further and possibly improve the extent of rRNA depletion. Nevertheless, as currently implemented, our method should enable the community to perform relatively easy, cost-effective, robust rRNA depletion, thereby facilitating RNA-seq studies.

## Materials and Methods

### Oligonucleotide algorithm

The algorithm was initialized with 500 and 1000 oligos of length 15 to 24 nucleotides for the 16S and 23S rRNA, respectively. Oligos were randomly positioned at non-gapped locations of the alignment of the 8 species we selected. Sequences were chosen by randomly selecting a nucleotide matching one or more species at each position. Sequences were then optimized to achieve the target predicted T_m_ of 62.5°C. T_m_ calculations were conducted using the MeltingTemp module in the biopython library. We used the default nearest-neighbor calculation table for RNA-DNA hybrids(12). Notably, this model does not allow prediction of T_m_ for some sequences with multiple sequential mismatches; as such, many oligos begin the optimization with undefined T_m_.

Optimization was conducted by sequential rounds of ‘mutation’ on each oligo. Allowed mutations included moving the probe from 1-4 bases, shrinking the probe from 1-4 bases (on either end), extending the probe from 1-4 bases (on either end), or swapping the sequence of the oligo at one position to a nucleotide matching a different aligned rRNA. In each round of mutation, the starting oligo was mutated 25 times. From this set of mutated oligos, an oligo close to the target T_m_ was chosen probabilistically (probabilities were determined by a normal distribution centered at 62.5°C with a standard deviation of 2°C). This probabilistic selection, coupled with the large number of oligos initialized, enables oligos to sample the possible binding locations without greedily descending on the first possible binding site they discover. Each oligo was mutated for 100 cycles before oligos binding to a number of sites across the 16S and 23S were selected.

To enable better binding of the more variable 23S 5’- and 3’-ends, we split the organisms into two groups (Ec, Pa, Cc, Rp and Ms, Mtb, Bs, Sa) and re-ran the optimization algorithm as above. For each of these groups, we selected 2 additional oligos matching the 5’- and 3’-ends of the 23S.

### Data and code availability

The code used to generate the oligonucleotides is available for download at https://github.com/peterculviner/ribodeplete. The raw and processed sequencing data is available on GEO (GSE142656).

### Bacterial strains and culture condition

*E. coli* MG1655 was grown to mid-log phase at 37 °C in LB medium or M9 medium supplemented with 0.1% casamino acids, 0.4% glucose, 2 mM MgSO_4_, and 0.1 mM CaCl_2_. *C. crescentus* CB15N/NA1000 was grown to mid-log phase in PYE medium at 30 °C. *B. subtilis* 168 was grown to mid-log phase at 37 °C in LB medium. For quantifying changes in expression from antibiotic treatment, cells were harvested 5 minutes after adding chloramphenicol or rifampicin at 50 μg/mL or 25 μg/mL, respectively.

### RNA extraction

*E. coli* RNA was harvested by mixing 1 mL of cells with 110 μL of ice-cold stop solution (95% ethanol and 5% acid-buffered phenol) and spinning in a table-top centrifuge for 30 s at 13000 rpm. *C. crescentus* RNA was harvested by spinning down 2 mL of cells in a table-top centrifuge for 30 s at 13000 rpm. After removing the supernatant, pellets were flash-frozen and stored at −80 °C until sample collection was complete. To extract RNA, TRIzol (Invitrogen) was heated to 65 °C and added to each cell pellet. The mixtures were then shaken at 65 °C for 10 min at 2000 rpm in a thermomixer and flash-frozen at −80 °C for at least 10 min. Pellets were thawed at room temperature and spun at top speed in a benchtop centrifuge at 4 °C for 5 min. The supernatant was added to 400 μL of 100% ethanol and passed through a DirectZol spin column (Zymo). Columns were washed twice with RNA PreWash buffer (Zymo) and once with RNA Wash buffer (Zymo), and RNA was eluted in 90 μL DEPC H_2_O. To remove genomic DNA, RNA was then treated with 4 μL of Turbo DNase I (Invitrogen) in 100 μL supplemented 10x Turbo DNase I buffer for 40 min at 37 °C. RNA was then diluted with 100 μL DEPC H_2_O, extracted with 200 μL buffered acid phenol-chloroform, and ethanol precipitated at −80 °C for 4 hr with 20 μL of 3 M NaOAc, 2 μL GlycoBlue (Invitrogen), and 600 μL ice-cold ethanol. Samples were centrifuged at 4 °C for 30 min at 21000 × *g* to pellet RNA, then washed twice with 500 μL of ice-cold 70% ethanol, followed by centrifugation at 4 °C for 5 min. RNA pellets were then air-dried and resuspended in DEPC H_2_O. RNA yield was quantified by a NanoDrop spectrophotometer, and RNA integrity was verified by running 50 ng of total RNA on a Novex 6% TBE-urea polyacrylamide gel (Invitrogen).

*B. subtilis* total RNA was harvested by mixing 5 mL of cell culture with 5 mL of cold (−30 °C) methanol and spinning down at 5000 rpm for 10 min. After removing the supernatant, pellets were frozen at −80 °C. To lyse cells, pellets were vortexted in 100 μL lysozyme (10 mg/mL) in TE (10 mM Tris-HCl and 1 mM EDTA) at pH = 8.0 and incubated for 5 min at 37 °C. Lysates were cleared by adding 350 μL Buffer RLT (Qiagen) in 1% beta-mercaptoethanol and vortexing. Lysates were then mixed with 250 μL ethanol, vortexed, and passed through an RNeasy mini spin column (Qiagen). Columns were washed with 350 μL Buffer RW1 (Qiagen). To remove genomic DNA, 40 μL of DNaseI in Buffer RDD (Qiagen) was applied to each column, and columns were incubated at room temperature for 15 min. Columns were then washed once with 350 μL Buffer RW1 (Qiagen) and twice with Buffer RPE (Qiagen), and RNA was eluted in 30 μL DEPC H_2_O. RNA yield was quantified by a NanoDrop spectrophotometer, and RNA integrity was verified by running 50 ng of total RNA on a Novex 6% TBE-urea polyacrylamide gel (Invitrogen).

### rRNA depletion, DIY method

Biotinylated oligos were selected using our algorithm, synthesized by IDT, and resuspended to 100 μM in Buffer TE (Qiagen). An undiluted oligo mix for each organism was created by mixing equal volumes of all 16S and 23S primers, as well as double volumes of 5S primers. This undiluted mix was then diluted based on the amount of total RNA added to the depletion reaction, using a custom bead calculator (available with code at https://github.com/peterculviner/ribodeplete).

Dynabeads MyOne Streptavidin C1 beads (ThermoFisher) were washed three times in an equal volume of 1x B&W buffer (5 mM Tris HCl pH = 7.0, 5 mM Tris HCl pH = 7.0, 500 μM EDTA, 1 M NaCl) and then resuspended in 30 μL of 2x B&W buffer (10 mM Tris HCl pH = 7.0, 10 mM Tris HCl pH = 7.0, 1 mM EDTA, 2 M NaCl). To prevent RNase contamination, 1 uL of SUPERAse-In RNase Inhibitor (ThermoFisher) was added to the beads. The beads were then incubated at room temperature until probe annealing (below) was complete.

To anneal biotinylated probes to rRNA, 2-3 μg total RNA, 20x SSC, 30 mM EDTA, water, and the diluted probe mix were mixed on ice in the calculated quantities. The mixtures were incubated in a thermocycler at 70 °C for 5 min, followed by a slow ramp down to 25 °C at a rate of 1 °C per 30 sec. To pull down biotinylated probes bound to rRNA, annealing reactions were then added directly to beads in 2x B&W buffer, mixed by pipetting and vortexing at medium speed, and incubated for 5 min at room temperature. Reactions were then vortexed on medium speed and incubated at 50 °C for 5 min, and then placed directly on a magnetic rack to separate beads from the remaining total RNA. The supernatant was pipetted away from the beads, placed on ice, and diluted to 200 μL in DEPC H_2_O. RNA was then ethanol precipitated at −20 °C for at least 1 hour with 20 μL of 3 M NaOAc, 2 μL GlycoBlue (Invitrogen), and 600 μL ice-cold ethanol. Samples were centrifuged at 4 °C for 30 min at 21000 × *g* to pellet RNA, then washed twice with 500 μL of ice-cold 70% ethanol, followed by centrifugation at 4 °C for 5 min. RNA pellets were then air-dried and resuspended in 10 μL DEPC H_2_O. RNA yield was quantified by a NanoDrop spectrophotometer, and the efficiency of rRNA depletion was verified by running 50 ng of total RNA on a Novex 6% TBE-urea polyacrylamide gel (Invitrogen).

### Optimization of rRNA depletion

In the process of generating our depletion protocol, we tried multiple ratios of streptavidin-coated beads to biotinylated oligos and biotinylated oligos to total RNA. We found that rRNA was depleted robustly across a range of ratios. However, it was critical to have a significant excess of streptavidin-coated beads over biotinylated oligos, as oligos that do not successfully capture rRNA may bind streptavidin more rapidly, thus out-competing bound rRNA-bound oligos and reducing rRNA capture efficiency. We selected our final ratios to achieve reliable depletion of rRNA at a low per-reaction cost.

### Cost calculation

The majority of reagents are common laboratory supplies for labs that work with RNA. To maintain the optimized ratio between streptavidin beads, biotinylated oligos, and rRNA, more oligos and beads must be used to deplete more total RNA. Considering the input, the cost per reaction is approximately $10, $19, or $28 for 1, 2 or 3 μg of RNA, respectively. The majority of the cost per reaction arises from streptavidin-coated magnetic beads; cost could likely be further decreased by using cheaper streptavidin-coated beads or decreasing the quantity of beads used (see above). The up-front cost of purchasing oligos (IDT) is approximately $1000 for large scale synthesis or $500 for smaller scale synthesis (available for sets of oligos >24). However, a single oligo synthesis order is adequate for hundreds of depletion reactions.

### RNA-seq library preparation

Libraries were generated as previously with a few modifications described below(13). The library generation protocol was a modified version of the paired-end strand-specific dUTP method using random hexamer priming. For libraries without rRNA removal, 500 ng of total RNA was used in the fragmentation step. For libraries with rRNA removal, 2-3 μg of input RNA was used in the rRNA removal step.

### rRNA depletion by Ribo-Zero

rRNA depletion via Ribo-Zero treatment (Illumina) was conducted as described previously(13). Briefly, provided magnetic beads were prepared individually by adding 225 μL of beads to a 1.5 mL tube, left to stand on a magnetic rack for 1 minute, washed twice with 225 μL of water, and resuspended in 65 μL of provided resuspension solution with 1 μL of provided RNase inhibitor. Samples were prepared using provided reagents with 4 μL of reaction buffer, 2-3 μg of total RNA, 10 μL of rRNA removal solution in a total reaction volume of 40 μL. Samples were incubated at 68 °C for 10 minutes and at room temperature for 5 minutes. Samples were added directly to the resuspended magnetic beads, mixed by pipetting, incubated for 5 minutes at room temperature, and then incubated for 5 minutes at 50 °C. After incubation, samples were placed on magnetic rack and the supernatant was transferred to a new tube, discarding the beads. Samples were ethanol precipitated as above with a 1 hour incubation at −20 °C and resuspended in 9 μL of water.

### Fragmentation

RNA libraries were fragmented by adding 1 μL of 10x fragmentation buffer (Invitrogen) to 9 μL of input RNA in DEPC H_2_O and heating at 70 °C for 8 min. Fragmentation reactions were stopped by immediately placing on ice and adding 1 μL of stop solution (Invitrogen). Reactions were diluted to 20 μL in DEPC H_2_O, and RNA was ethanol precipitated at −20 °C for at least 1 hour with 2 μL of 3 M NaOAc, 2 μL GlycoBlue (Invitrogen), and 60 μL ice-cold ethanol. Samples were centrifuged at 4 °C for 30 min at 21000 × *g* to pellet RNA, then washed with 200 μL of ice-cold 70% ethanol, followed by centrifugation at 4 °C for 5 min. RNA pellets were then air-dried and resuspended in 6 μL DEPC H_2_O.

### cDNA synthesis

1 μL of random primers at 3 μg/μL (Invitrogen) were added to fragmented RNA, and the mixture was heated at 65 °C for 5 min and placed on ice for 1 min. To conduct first strand synthesis, 4 μL of first strand synthesis buffer (Invitrogen), 2 μL of 100 mM DTT, 1 μL of 10 mM dNTPs, 1 μL of SUPERase-In (Invitrogen), and 4 μL of DEPC H_2_O were added to each reaction. Reaction mixtures incubated at room temperature for 2 minutes, followed by addition of 1 μL of Superscript III. Reactions were then placed in a thermocycler for the following program: 25 °C for 10 min, 50 °C for 1 hr, and 70 °C for 15 min. To extract cDNA, reactions were diluted to 200 μL in DEPC H_2_O, then vortexed with 200 μL of neutral phenol-chloroform isoamyl alcohol. Following centrifugation, the aqueous layer was extracted, and cDNA was ethanol precipitated at −20 °C for at least 1 hour with 18.5 μL of 3 M NaOAc, 2 μL GlycoBlue (Invitrogen), and 600 μL ice-cold ethanol. Samples were centrifuged at 4 °C for 30 min at 21000 × *g* to pellet cDNA, then washed twice with 500 μL of ice-cold 70% ethanol, followed by centrifugation at 4 °C for 5 min. Pellets were then air-dried and resuspended in 104 μL DEPC H_2_O. Second strand synthesis was conducted by adding 30 μL of second strand synthesis buffer (Invitrogen), 4 μL of 10 mM dNTPs (with dUTP instead of dTTP), 4 μL of first strand synthesis buffer (Invitrogen), and 2 μL of 100 mM DTT to each sample, followed by incubation on ice for 5 min. To initiate second strand synthesis, 1 μL of RNase H (NEB), 1 μL of *E. coli* DNA ligase (NEB), and 4 μL of *E. coli* DNA polymerase I (NEB) were added to each sample. Reactions were then incubated at 16 °C for 2.5 hr.

### End-repair and adaptor ligation

Cleanup for second strand synthesis and all subsequent steps was conducted using Agencourt AMPure XP magnetic beads (Beckman Coulter), and beads were left in the reaction to be reused for subsequent cleanup steps. For each sample, 100 μL of beads were added to 1.5 mL tubes and placed on a magnetic rack. The supernatant was removed and replaced with 450 μL of 20% (w/v) PEG 8000 in 2.5 M NaCl. Second strand synthesis reactions were then added directly to resuspended beads, mixed by pipetting and vortexing, and incubated at room temperature for 5 min. Samples were then placed on a magnetic rack for ~10 min, or until the solution was clear, and the supernatant was removed. Beads were then washed twice in 500 μL of 80% ethanol, dried, and resuspended in 50 μL of elution buffer (Qiagen). End repair reactions were conducted by adding 10 μL of 10x T4 DNA ligase buffer (NEB), 4 μL of 10 mM dNTPs, 5 μL of T4 DNA polymerase (NEB), 1 μL of Klenow DNA polymerase (NEB), 5 μL of T4 polynucleotide kinase (NEB), and 25 μL of DEPC H_2_O and incubating at 25 °C for 30 min. To clean up the reactions, 300 μL of 20% (w/v) PEG 8000 in 2.5 M NaCl was mixed with each reaction by pipetting and vortexing. Samples were then incubated at room temperature for 5 min, and then placed on a magnetic rack for ~5 min. The supernatant was removed, and the beads were then washed twice in 500 μL of 80% ethanol, dried, and resuspended in 32 μL of elution buffer (Qiagen). 3’-adenylation reactions were conducted by adding 5 μL of NEB buffer 2 (NEB), 1 μL 10 mM dATP, 3 μL Klenow fragment (3’→5’ exo-) (NEB), and 9 μL of DEPC H_2_O to each reaction and incubating at 37 °C for 30 min. To clean up the reactions, 150 μL of 20% (w/v) PEG 8000 in 2.5 M NaCl was mixed with each reaction by pipetting and vortexing. Samples were then incubated at room temperature for 5 min, and then placed on a magnetic rack for ~5 min. The supernatant was removed, and the beads were then washed twice in 500 μL of 80% ethanol, dried, and resuspended in 20 μL of elution buffer (Qiagen). To elute DNA from the beads, reactions were incubated at room temperature for 5 min. Tubes were then returned to the magnetic rack and incubated for 1-2 min to allow the solution to clear, and then half of the supernatant (10 μL) was removed and stored at −20 °C in case of downstream failure. To ligate adaptors to DNA, 1 μL of 5 μM annealed adaptors and 10 μL of Blunt/TA ligase master mix (NEB) was added to each reaction, and reactions were incubated at 25 °C for 20 min. Annealed adaptor mix was made by mixing 25 μL of a 200 μM solution of each paired-end adaptor together, heating to 90°C for 2 minutes, cooling at 2°C/minute for 30 minutes on a thermocycler, placing on ice, adding 50 μL of water, and storing aliquots at −20°C. To clean up ligation reactions, 60 μL of 20% (w/v) PEG 8000 in 2.5 M NaCl was mixed with each reaction by pipetting and vortexing, and reactions were incubated at room temperature for 5 min. Reactions were then placed on a magnetic rack for ~10 min, until solutions were clear, and the supernatant was removed. The beads were then washed twice in 500 μL of 80% ethanol, dried, and resuspended in 19 μL of 10 mM Tris-HCl (pH = 8) and 0.1 mM EDTA. Reactions were then incubated at room temperature for 5 min to completely elute DNA. Tubes were then returned to the magnetic rack and incubated for 1-2 min to allow the solution to clear, and then the supernatant was removed and moved to a new tube and the beads discarded. To digest the dUTP-containing second strand, 1 μL of USER enzyme (NEB) was added to 19 μL of eluted DNA and incubated at 37 °C for 15 min, followed by heat-inactivation at 95 °C for 5 min.

### Library amplification

PCR reactions were prepared by mixing 10 μL of library template (diluted if too concentrated), 2 μL of 25 μM global primer, 2 μL of 25 μM barcoded primer, 11 μL of H_2_O, and 25 μL of 2x KAPA HiFi HotStart ReadyMix (Roche). Reactions were then cycled through the following thermocycler protocol: 98 °C/45 s, 98 °C/15 s, 60 °C/30 s, 72 °C/30 s, 72 °C/1 min. Steps 2-4 were repeated for 9-12 cycles, depending on the results of 10 μL optimization reactions. Following amplification, PCR reactions were run on an 8% TBE polyacrylamide gel (Invitrogen) for 30 min at 180 V, and the region from 200 to 350 bp was excised, crushed, soaked in 500 μL 10 mM Tris pH = 8.0, and frozen at −20 °C for at least 15 min. To elute DNA from the gel, reactions were shaken at 2000 rpm for 10 min at 70 °C in a thermomixer, followed by 1 hr at 37 °C. Reactions were then spun through a Spin-X 0.22 μm cellulose acetate column (Costar) and transferred to a new tube. Libraries were isopropanol precipitated by adding 32 μL 5 M NaCl, 2 μL GlycoBlue (Invitrogen), and 550 μL 100% isopropanol and incubating at −20 °C for at least 1 hr. Samples were then centrifuged at 4 °C for 30 min at 21000 × *g* to pellet DNA, then washed with 1 mL of ice-cold 70% ethanol, followed by centrifugation at 4 °C for 5 min. DNA pellets were then air-dried and resuspended in 11 μL H_2_O. Paired-end sequencing of amplified libraries was then performed on an Illumina NextSeq500, and single-end sequencing on an Illumina MiSeq.

### RNA-sequencing read mapping and normalization

FASTQ files for each barcode were mapped to the *E. coli* MG1655 genome (NC_000913.2), the *B. subtilis* 168 genome (NC_000964.3), or the *C. crescentus* NA1000 genome (NC_011916.1) using bowtie2 (version 2.1.0) with the following arguments: -D 20 -R 3 -N 0 -L 20 -i S,1,0.50. The samtools (version 0.1.19) suite was used via the pysam library (version 0.9.1.4) for interconversion of BAM and SAM file formats and conducting indexing. Gene names and coding region positions were extracted from NCBI annotations.

### Single-end sequencing

For all analyses except that of fragment density across *E. coli* rRNA loci, one count was added to the middle of each read. All reads mapping to a given coding region were then summed and normalized by reads per kilobase of transcript per million (RPKM). This normalized quantity was then used in all downstream analyses.

For analysis of fragment density across rRNA loci, one count was added for all positions between and including the 5’-and 3’-ends of reads. To correct for variability in sequencing depth, counts at each position were divided by a sample size factor. Briefly, counts recorded in each genomic region were summed for all samples and then the geometric mean was taken across samples to yield a reference sample. The size factor for a given sample was the median counts in all regions after normalizing counts to the reference samples.

### Analysis of oligo depletion efficiency

To quantify the efficiency of rRNA depletion, the sum of reads mapping to rRNA loci was divided by the total number of mapped reads in each sample. To compare the reads mapping to individual coding regions following rRNA depletion and/or antibiotic treatment (Figures 3A-C, S2A-F, and S3A), coding regions were filtered for expression by RPKM, and then the correlation between RPKM for individual coding regions was compared using the SciPy statistical functions package. Outliers for the ratio of reads per coding region following Ribo-Zero versus DIY treatment (Figure S2A) were identified by measuring the distance for all genes in Cartesian coordinates from the log-log least squares fit for all regions above the expression threshold. Outliers were defined as genes for which this ratio was less than or greater than two standard deviations from the mean line. Outliers in Figure S2D were hand-picked.

## Acknowledgements

The co-first authors discussed and mutually agreed on the author order. We thank I. Frumkin and I. Nocedal for comments on the manuscript, D. Parker and M. Guzzo for reagents, and M. Guo for helpful discussions. This work was funded by an NIH grant to M.T.L. (R01GM082899), who is also an Investigator of the Howard Hughes Medical Institute. This work was also supported by NSF predoctoral graduate research fellowships to P.H.C. and C.K.G.

---

**Figure S1.**
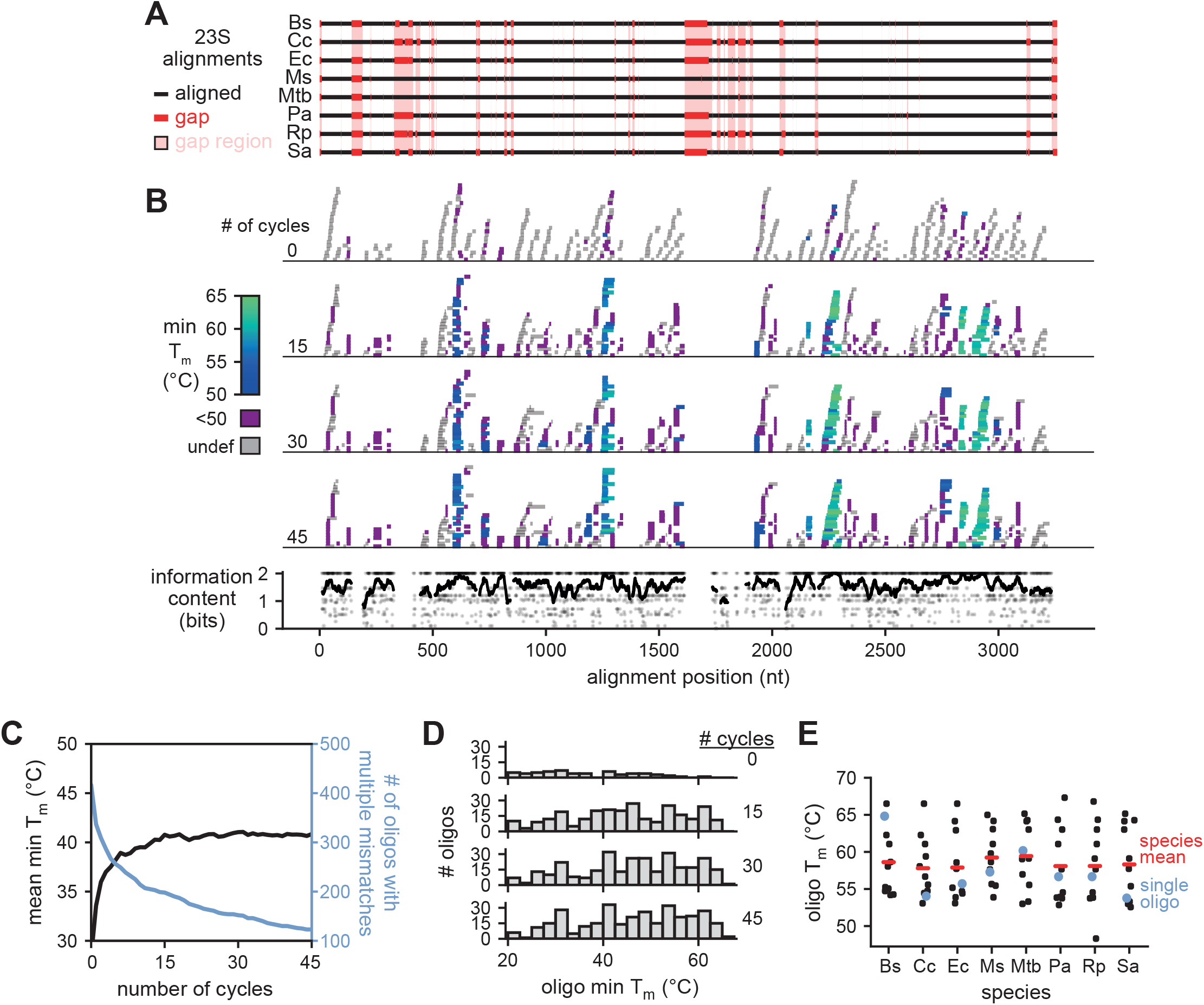
Oligonucleotide selection for 23S rRNA. (A) Alignments of all 23S sequences from 8 bacterial species (Ec = *E. coli*; Pa = *P. aeruginosa*; Rp = *R. parkeri*; Cc = *C. crescentus*; Bs = *B. subtilis*; Ms = *M. smegmatis*; Mtb = *M. tuberculosis*; Sa = *S. aureus*). Alignment gaps are shown as red lines in the particular species of the gap. Regions with a gap in any species are highlighted in pink; these regions were not considered when designing oligos. (B) The position, length, and minimum T_m_ of all oligos plotted against the 23S alignment after the indicated number of optimization cycles (top). The information content at each nucleotide position of aligned regions is also shown (bottom, points). To highlight conserved regions, a sliding average information content is also plotted (bottom, line). (C) Oligo T_m_ statistics after multiple cycles of the T_m_ optimization algorithm. For each oligo (n = 500), we calculated the minimum T_m_ across the 8 species considered and then plotted the mean of this value across all oligos (black). The T_m_ cannot be accurately estimated for oligos with multiple sequential mismatches; the number of oligos with an undefined T_m_ is also plotted (blue). (D) Histograms of minimum T_m_ for oligos at the indicated number of optimization cycles. Data were generated as in (C), but oligo T_m_ minima were used to generate histograms rather than taking the mean across all oligos. Oligos with undefined T_m_ were not included in the histograms. (E) Distribution of T_m_ values for each 23S-targeting oligo (n = 11) for each individual species indicated. The mean T_m_ of oligos for each species is also shown (red lines). Note that the same oligos are used for each species, but because of 23S sequence variability, the T_m_ can vary, as illustrated for one particular oligo (blue).

**Figure S2.**
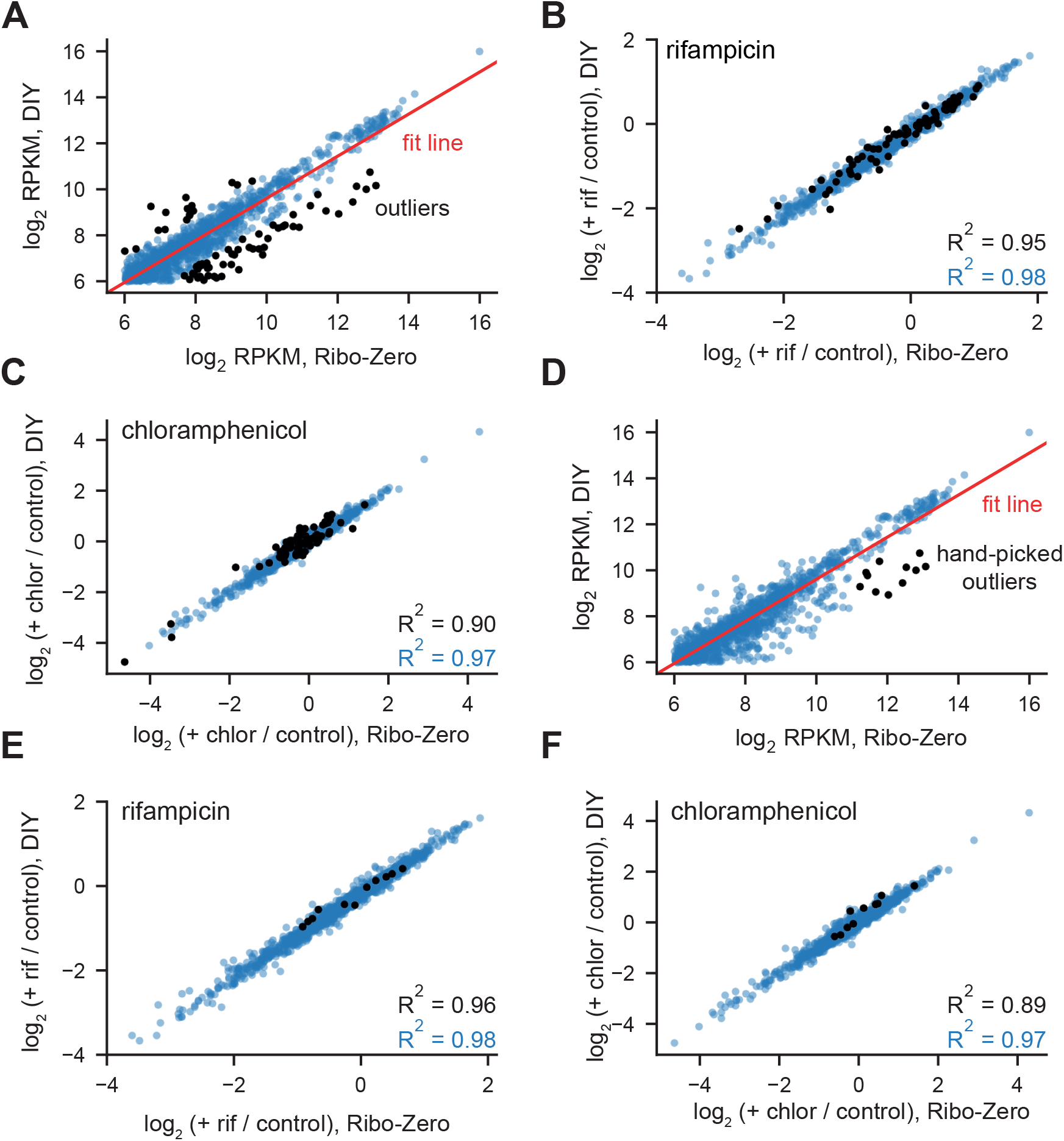
Analysis of outliers in correlation between mRNA counts following Ribo-Zero and DIY rRNA depletion. (A) Figure 3C, with all genes at least two standard deviations away from the least squares fit line (red) indicated in black (n = 69). (B) Figure 3A, with outliers identified in Figure S3A marked in black. For these outliers, the correlation between log_2_ (rif+/negative control) for DIY depletion and Ribo-Zero treatment was 0.95, compared to 0.98 for all well-expressed coding regions. (C) Figure 3B, with outliers identified in Figure S3A marked in black. For these outliers, the correlation between log_2_ (chl+/negative control) for DIY depletion and Ribo-Zero treatment was 0.90, compared to 0.97 for all well-expressed coding regions. (D) Figure 3C, with 11 highly-expressed genes more depleted in our method than in Ribo-Zero indicated in black. (E) Figure 3A, with outliers identified in Figure S3D marked in black. For these outliers, the correlation between log_2_ (rif+/negative control) for DIY depletion and Ribo-Zero treatment was 0.96, compared to 0.98 for all well-expressed coding regions. (F) Figure 3B, with outliers identified in Figure S3D marked in black. For these outliers, the correlation between log_2_ (chl+/negative control) for DIY depletion and Ribo-Zero treatment was 0.89, compared to 0.97 for all well-expressed coding regions.

**Figure S3.**
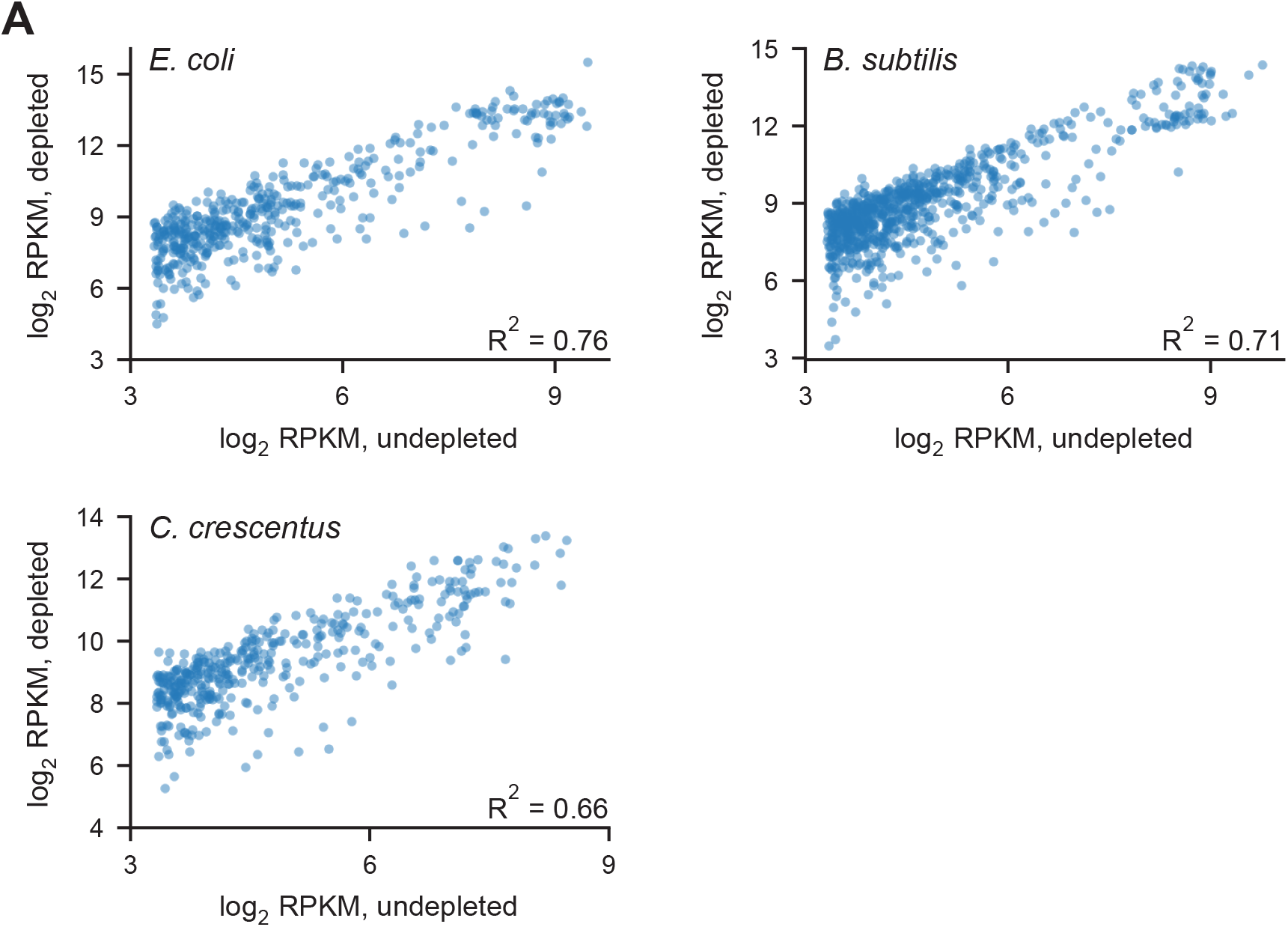
Correlation between counts per coding region pre- and post-rRNA depletion for *B. subtilis* and *C. crescentus* total RNA. (A) Top left: scatterplot showing correlation between read counts (RPKM) for *E. coli* coding regions pre- and post-rRNA depletion using our depletion strategy. All coding regions with at least 10 counts in both samples (n = 438) were considered in this analysis. Top right: scatterplot showing correlation between read counts (RPKM) for *B. subtilis* coding regions pre- and post-rRNA depletion using our depletion strategy. All coding regions with at least 10 counts in both samples (784 regions total) were considered in the analysis. Bottom: scatterplot showing correlation between read counts (RPKM) for *C. crescentus* coding regions pre- and post-rRNA depletion using our depletion strategy. All coding regions with at least 10 counts in both samples (398 regions total) were considered in the analysis. (B) Fold-depletion for various ratios of oligo probe : RNA and streptavidin bead : oligo probe ratios. Depletions were calculated by qRT-PCR to a single region within each rRNA relative to a bead-only negative control.

**Table S1.**
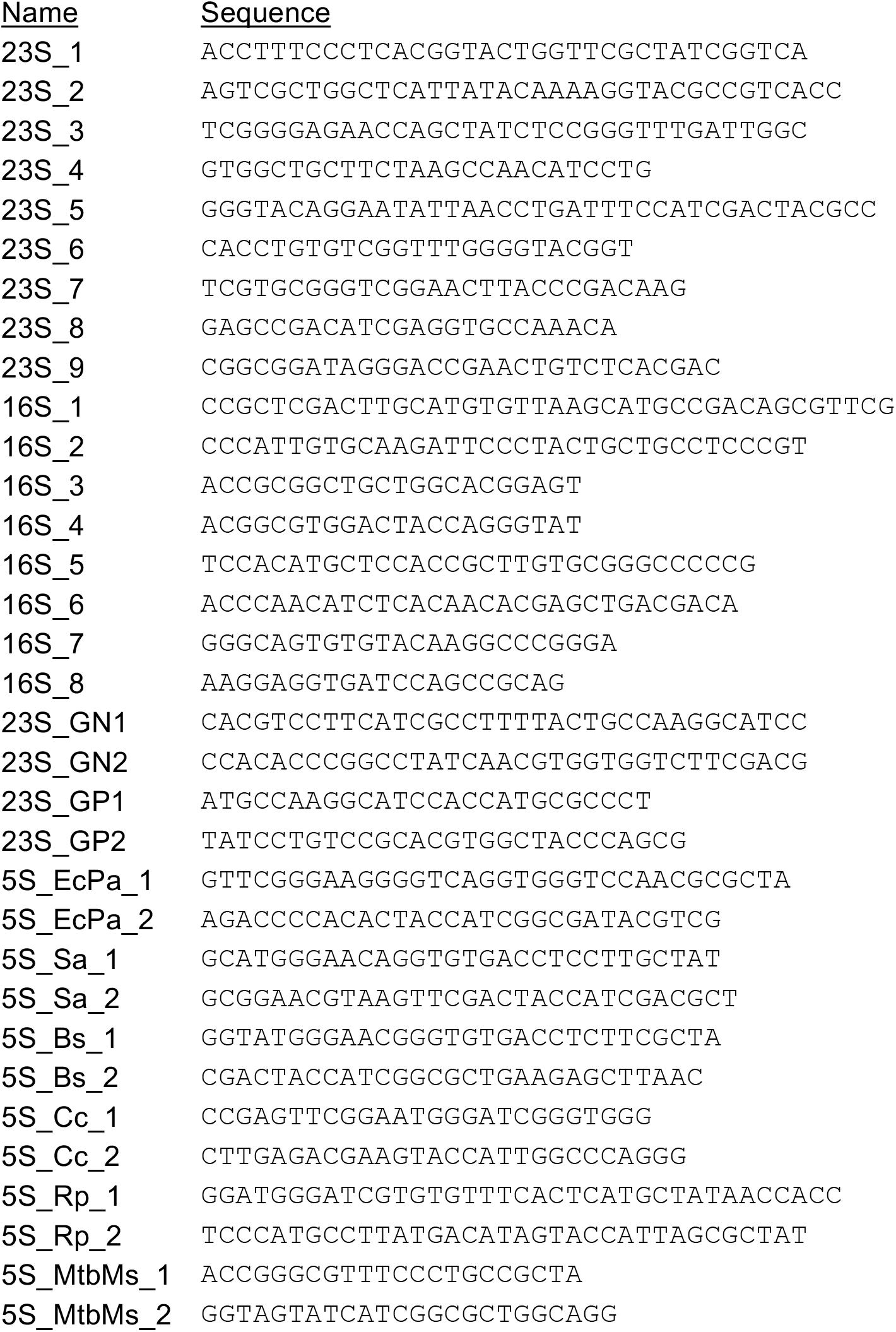
Sequences of oligonucleotides for bacterial rRNA depletion

## References

1. Hör J, Gorski SA, Vogel J. 2018. Bacterial RNA Biology on a Genome Scale. Mol Cell 70:785–799.

2. Croucher NJ, Thomson NR. 2010. Studying bacterial transcriptomes using RNA-seq. Curr Opin Microbiol 13:619–624.

3. Creecy JP, Conway T. 2015. Quantitative bacterial transcriptomics with RNA-seq. Curr Opin Microbiol 23:133–140.

4. Podnar J, Deiderick H, Huerta G, Hunicke-Smith S. 2014. Next-generation sequencing RNA-Seq library construction. Curr Protoc Mol Biol 1–19.

5. Nagalakshmi U, Wang Z, Waern K, Shou C, Raha D, Gerstein M, Snyder M. 2008. The transcriptional landscape of the yeast genome defined by RNA sequencing. Science (80-) 320:1344–1349.

6. Mortazavi A, Williams BA, McCue K, Schaeffer L, Wold B. 2008. Mapping and quantifying mammalian transcriptomes by RNA-Seq. Nat Methods 5:621–628.

7. Westermann AJ, Gorski SA, Vogel J. 2012. Dual RNA-seq of pathogen and host. Nat Rev Microbiol 10:618–630.

8. Herbert ZT, Kershner JP, Butty VL, Thimmapuram J, Choudhari S, Alekseyev YO, Fan J, Podnar JW, Wilcox E, Gipson J, Gillaspy A, Jepsen K, BonDurant SS, Morris K, Berkeley M, LeClerc A, Simpson SD, Sommerville G, Grimmett L, Adams M, Levine SS. 2018. Cross-site comparison of ribosomal depletion kits for Illumina RNAseq library construction. BMC Genomics 19:1–10.

9. Stewart FJ, Ottesen EA, Delong EF. 2010. Development and quantitative analyses of a universal rRNA-subtraction protocol for microbial metatranscriptomics. ISME J 4:896–907.

10. He S, Wurtzel O, Singh K, Froula JL, Yilmaz S, Tringe SG, Wang Z, Chen F, Lindquist EA, Sorek R, Hugenholtz P. 2010. Validation of two ribosomal RNA removal methods for microbial metatranscriptomics. Nat Methods 7:807–812.

11. Huang Y, Sheth RU, Kaufman A, Wang HH. 2019. Scalable and cost-effective ribonuclease-based rRNA depletion for transcriptomics. Nucleic Acids Res.

12. Sugimoto N, Nakano S, Katoh M, Matsumura A, Nakamuta H, Ohmichi T, Yoneyama M, Sasaki M. 1995. Thermodynamic Parameters To Predict Stability of RNA/DNA Hybrid Duplexes. Biochemistry 34:11211–11216.

13. Culviner PH, Laub MT. 2018. Global Analysis of the E. coli Toxin MazF Reveals Widespread Cleavage of mRNA and the Inhibition of rRNA Maturation and Ribosome Biogenesis. Mol Cell 70:868–880.e10.

